# A Far-Red Fluorescent Probe to Visualize *Staphylococcus aureus* in Patient Samples

**DOI:** 10.1101/2023.09.04.556223

**Authors:** Krittapas Jantarug, Vishwachi Tripathi, Benedict Morin, Aya Iizuka, Richard Kuehl, Mario Morgenstern, Martin Clauss, Nina Khanna, Dirk Bumann, Pablo Rivera-Fuentes

## Abstract

*Staphylococcus aureus (S. aureus)* is the leading bacterial cause of death in high-income countries and can cause invasive infections at various body sites. These infections are associated with prolonged hospital stays, a large economic burden, considerable treatment failure, and mortality rates. So far, there is only limited knowledge about the specific locations where *S. aureus* resides in the human body during various infections. Hence, the visualization of *S. aureus* holds significant importance in microbiological research. Herein, we report the development and validation of a far-red-fluorescent probe to detect *S. aureus* in human biopsies from deep-seated infections. This probe displays strong fluorescence and low background in human tissues, outperforming current tools for *S. aureus* detection. Several applications are demonstrated, including fixed- and live-cell imaging, flow cytometry, and super-resolution bacterial imaging.

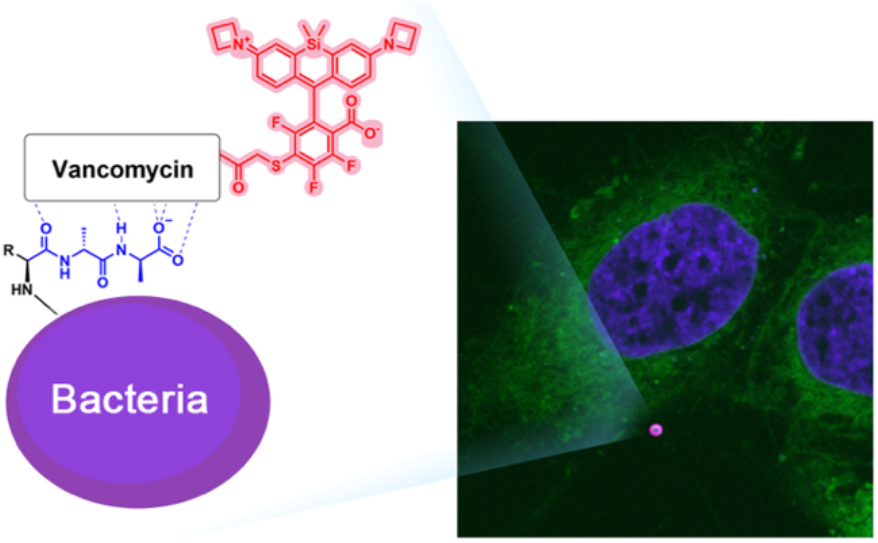

## Introduction

*Staphylococcus aureus (S. aureus)* stands as the primary bacterial contributor to death in wealthy countries.^1^ *S. aureus* commonly infects the skin and soft tissues but can also spread directly or via the bloodstream to virtually any site of the body including bone and joints, lungs, heart valves or central nervous system.^2,3^ These infections are associated with prolonged hospital stays, a large economic burden, 20-30% treatment failure, and mortality rates of up to 20-40%.^4,5^ *S. aureus* infections are often resilient to antibiotic treatment, even if the causative *S. aureus* strain appears susceptible to the employed antibiotics in clinical microbiology laboratories. These infections often need mechanical removal of the primary site of infection by surgery or drainage for a successful treatment. To date, local conditions and mechanisms in infected human tissues remain poorly characterized. Thus, a better understanding of the pathological mechanisms underlying the persistence of *S. aureus* in human tissue is necessary.

Fluorescent tools have been developed to detect *S. aureus* in the context of mammalian cells and tissues. Immunostaining is frequently used to label bacteria of interest in tissue sections. An antibody against *S. aureus* surface proteins (anti-SA) is commonly applied to detect *S. aureus*,^6^ but exhibits staining heterogeneity due to varying levels of protein expression. An antibody against wall teichoic acids (anti-WTA) detects *S. aureus* more comprehensively but is expensive and synthesis is protected by patent.^7,8^ Alternatively, antibiotics conjugated with fluorophores can be used as bacterial detection tools.^9^ These small-molecule probes can have significant advantages over antibodies, including better tissue penetration, higher chemical stability, compatibility with antibody-based detection of other biomolecules, and lower cost. This kind of probe has been developed to image *S. aureus* by leveraging the binding of the glycopeptide vancomycin to D-alanyl-D-alanine moieties of lipid II (a precursor of peptidoglycan) and the mature peptidoglycan layer of bacteria.^10^ Vancomycin labeled with the green fluorophore boron dipyrromethene (Van-BODIPY)^11^ or with a near-infrared dye (Vanco-800CW) have been reported (Figure S1).^12^ These probes, however, also exhibit drawbacks such as spectral overlap with host autofluorescence, which is particularly high in inflamed tissues (Van-BODIPY),^11,13^ or an excitation wavelength that is not commonly available in fluorescence microscopes or flow cytometers (Vanco-800CW).

## Results

We posited that detection of *S. aureus* in mammalian cells and human tissues, including patient biopsies, would benefit from a different fluorophore that is bright and photostable, has minimal fluorescence background regardless of the sample heterogeneity or preservation method, is simple to use, and can be analyzed by common instruments found in research labs. Considering these challenges, we selected the JF_669_ dye as the fluorescent reporter.^14^ This fluorophore is bright and photostable at a long wavelength, yet it can be excited with a common laser used in microscopes and flow cytometers. Moreover, compared to the popular JF_646_ dye,^15^ JF_669_ is more photostable and is more fluorogenic,^14^ i.e., it produces a brighter signal upon binding to an intended macromolecular target.

Probe Van-JF_669_ was conveniently prepared in one step from commercially available vancomycin and JF_669_-NHS ester (Figure 1A). Structural analysis by high-resolution tandem mass spectrometry revealed that conjugation of the dye occurs selectively at the primary amine on the sugar moiety of vancomycin (Figure S2), thus this synthesis produces a homogenous probe with a well-defined structure. Moreover, the primary amine on the disaccharide does not have specific interactions with D-alanyl-D-alanine,^16^ thus installation of the fluorophore at this position does not prevent binding to peptidoglycan.

**Figure 1.**
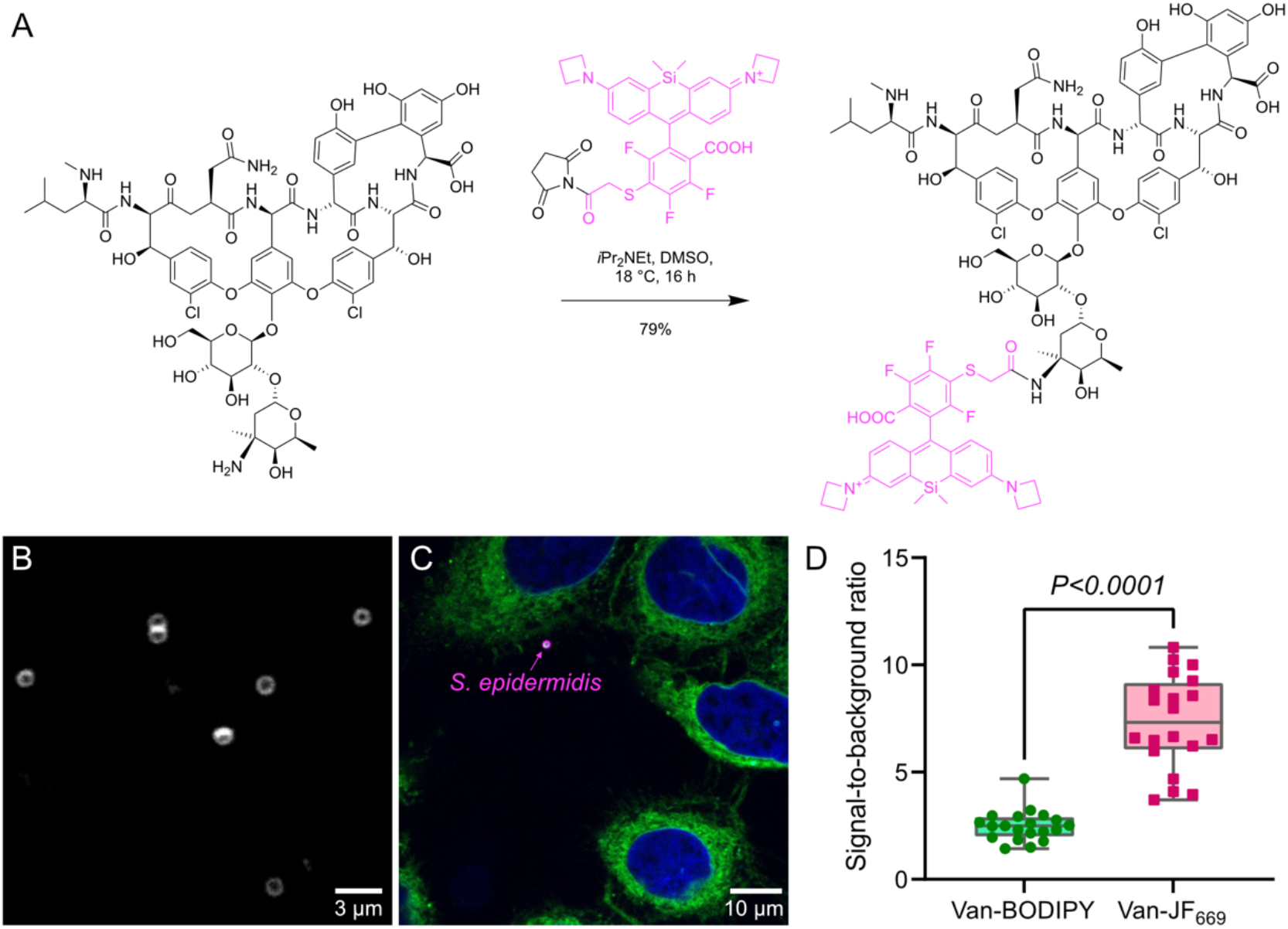
Synthesis and in vitro validation of Van-JF_669_. (A) Synthesis of Van-JF_669_. (B) Micrograph of *S. epidermidis* treated with 2 nM of Van-JF_669_. (C) Micrograph of *S. epidermidis* co-cultured with HeLa cells treated with 1.5 μM Hoechst (nuclei = blue), 2 μM ER-Tracker™ Green (endoplasmic reticulum = green) and 10 nM Van-JF_669_ (*S. epidermidis* = magenta). (D) Signal-to-background ratio of *S. epidermidis* co-cultured with HeLa cells labeled Van-BODIPY or Van-JF_669_. *N* = 20 independent cells per sample were examined. Statistical significance was evaluated by paired *t*-test. The horizontal line inside the box indicates the median, the box indicates the mean ± standard deviation, and whiskers indicate the minimum and maximum values.

The excitation and emission spectra of Van-JF_669_ displayed essentially the same wavelengths as those of the free dye, and the quantum yield also remained unchanged (ϕ_JF669_ = 0.37,^14^ ϕ_Van-JF669_ = 0.37). Van-JF_669_ preserved the polarity-dependent fluorogenicity of the parent dye, displaying low fluorescence in apolar medium (Figure S3). This behavior contrasts with those of traditional dyes, such as BODIPY, which have stronger fluorescence in media of low polarity (Figure S3) and thus display large background fluorescence due to binding to hydrophobic substances such as membranes. Van-JF_669_ exhibited a minimum inhibitory concentration (MIC) of 5 mg L^−1^ (2.3 μM) against lab strain methicillin-resistant *S. aureus* (MRSA, ATCC 43300), lab strain methicillin-sensitive *S. aureus* (MSSA, ATCC 29213), and clinical isolate *S. aureus* (MSSA, PROSA28), which is significantly higher than vancomycin (MIC = 0.31 mg L^−1^ - 0.2 μM) in the same strains (Figure S4). This MIC difference indicates that the attached fluorophore diminished vancomycin’s affinity for its target D-alanyl-D-alanine, similar to other conjugates attached at the primary amine.^10,16^ The lower toxicity of Van-JF_669_ compared to the parent antibiotic is an advantage since it allows for imaging live cells with limited impact on bacterial viability.

We first evaluated the labeling efficiency of Van-JF_669_ with *Staphylococcus epidermidis* (*S. epidermidis*, ATCC 12228). *S. epidermidis* were treated with Van-JF_669_ (2 nM) and images were acquired with a spinning disk confocal fluorescence microscope without any washing step. Strong fluorescence was observed on the cell wall of *S. epidermidis* (Figure 1B). Van-JF_669_ at 10 nM specifically stained *S. epidermidis* cells but not human cells (Figure 1C and Figure S5). This high selectivity permitted imaging without a necessity to wash away the staining solution. Van-BODIPY also labeled the bacterial cell wall (Figure S5), but the signal-to-background ratio was significantly lower than that of Van-JF_669_ (Figure 1D).

Van-JF_699_ also selectively stained *S. aureus* in cell-culture infections of human monocyte-like THP-1 cells (ATCC TIB-202) with *S. aureus* (Cowan). Imaging flow cytometry revealed selective staining of *S. aureus*. The signal-to-background was higher compared to an alternative labeling method based on cell-wall incorporation of a tetramethyl rhodamine-functionalized D-amino acid (RADA) ^18^ (Fig. 2A-C). Van-JF_669_ also stained the clinical isolate *S. aureus* (PROSA25) in primary monocyte-derived macrophages isolated from healthy human donors (Fig. 2D).

**Figure 2.**
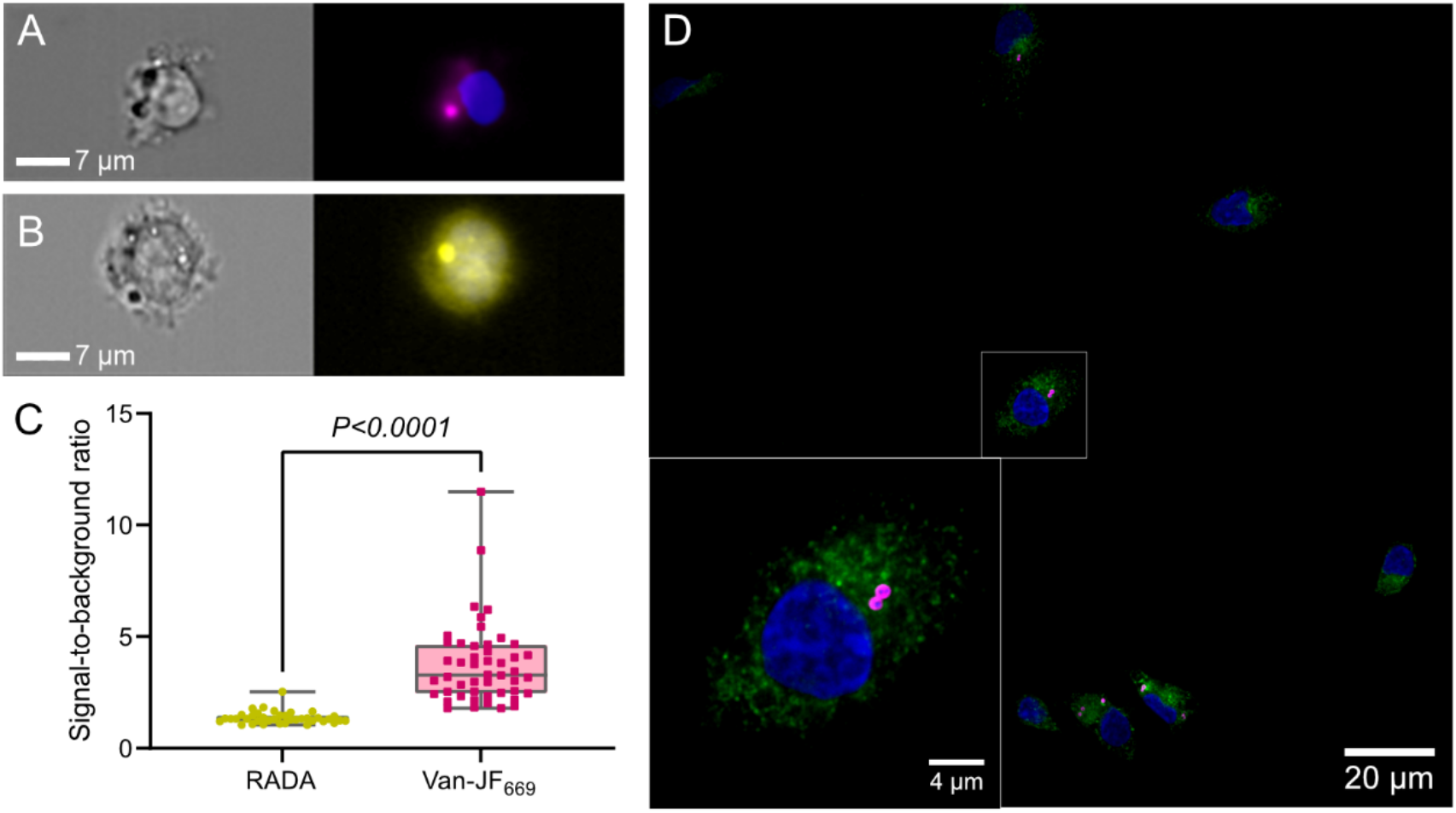
Flow cytometry image of in vitro co-culture of THP1 cells with clinical *S. aureus* (PROSA25) labeled with (A) 1.2 μM Van-JF_669_ and (B) 4.84 μM RADA treated with 3 μM DAPI (nuclei = blue). (C) Signal-to-background ratio of *S. aureus* in THP1 cells labeled RADA and Van-JF_669_. *N* = 38 and 48 independent cells per time point were examined (from left to right). Statistical significance was evaluated by paired t-test. The horizontal line inside the box indicates the median, the box indicates the mean ± standard deviation, and whiskers indicate the minimum and maximum values. (D) Fluorescent images of clinical *S. aureus* (PROSA25)-primary macrophages co-cultured. The clinical *S. aureus* was phagocyted by primary macrophages followed by treatment of 240 nM Van-JF_669_(*S. aureus* = magenta), anti-human Rab5 (human cells = green), and 3 μM DAPI (nuclei = blue). The inset displays a magnified view of the area marked with a white rectangle in the main image.

Van-JF_669_ also stained a clinical isolate of *S. aureus* (PROSA28). To enhance staining of live *S. aureus*, we mixed 130 nM Van-JF_669_ with 170 nM unlabeled vancomycin, which promotes the formation of heterodimers of these compounds with increased affinity for bacterial cell wall, particularly for live *S. aureus* (Supporting Video 1).^16,17^ *S. aureus* was able to divide under these conditions and the no-wash nature of the staining enabled efficient and sustained labeling of newly formed daughter cells.

To evaluate the labeling efficiency of Van-JF_669_ in patient tissues, we co-stained a biopsy section from a patient with osteomyelitis using Van-JF_669_ and an antibody to *S. aureus* protein A (anti-SA). We observed optimal staining of *S. aureus* cell wall in these samples at slightly higher concentrations (130 nM), probably because of low-level adsorption to the tissue. Whereas anti-SA stained only the outer surface protein of *S. aureus*, Van-JF_669_ stained also the septum between two emerging daughter cells, which is an area of cell wall with particularly active de novo peptidoglycan synthesis (Fig. 3A-C).^19^ These data confirm the identity of Van-JF_669_-positive particles as *S. aureus*.

**Figure 3.**
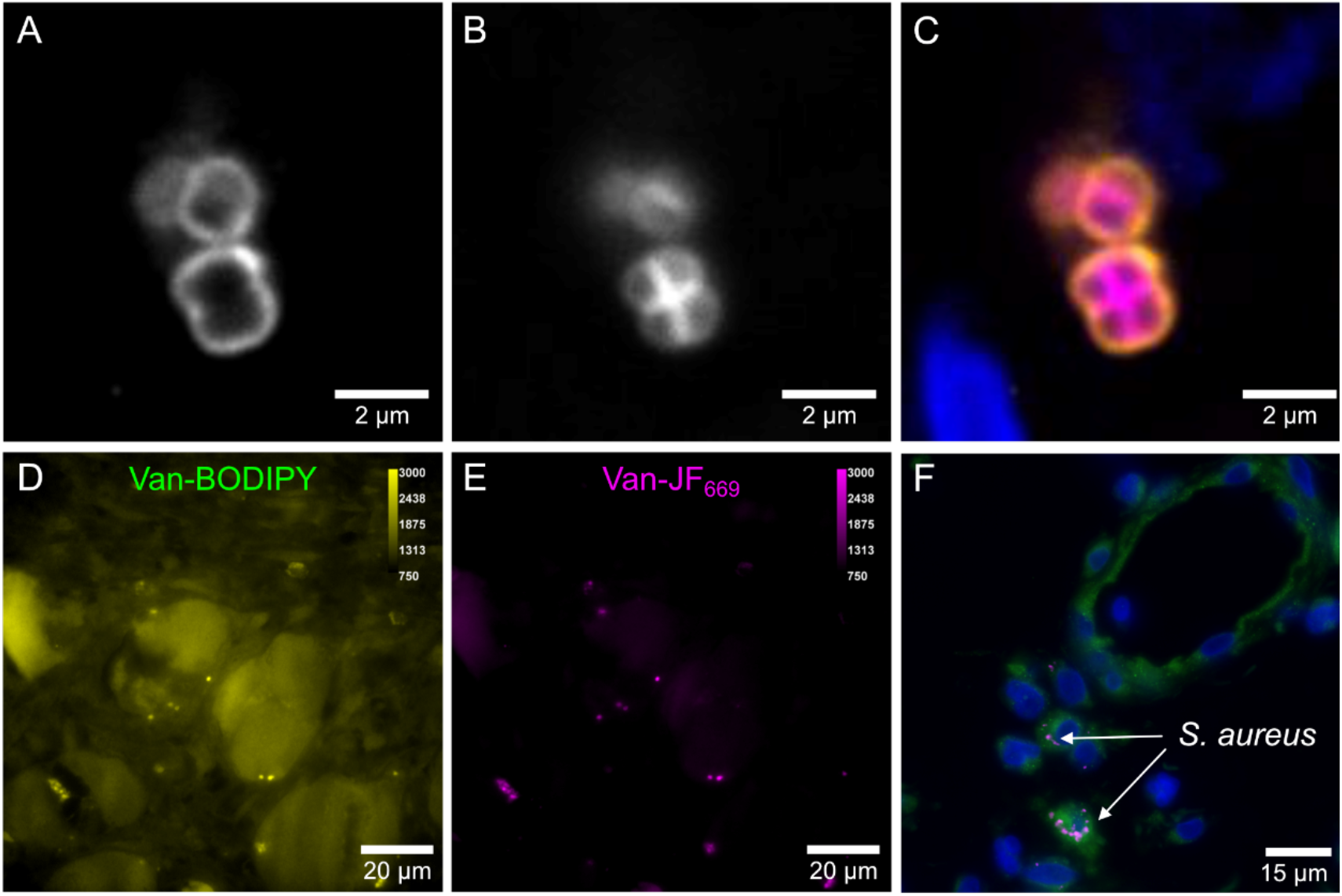
Fluorescence microscopy images of patient samples treated with fluorescent probes or antibodies. (A) 200 nM anti-SA-AF568, followed by (B) 130 nM Van-JF_669_ and 3 μM DAPI. (C) Merge of panels A and B (anti-SA-AF569 = orange, Van-JF_669_ = magenta, nuclei = blue). Fluorescent images of an FFPE-preserved patient sample with an *S. aureus* prosthetic joint infection stained with (D) 200 nM Van-BODIPY and (E) 130 nM Van-JF_669_. Color bars indicate pixel intensity values. (F) Multicolor fluorescent image of a biopsy from a patient with *S. aureus* prosthetic joint infection preserved by FFPE. The sample was treated with 130 nM Van-JF_669_ (*S. aureus* = white), anti-human Rab5 (human intracellular protein = green), and 3 μM DAPI (nuclei = blue).

In a formalin-fixed, paraffin-embedded (FFPE) section of a biopsy from a prosthetic joint infection, staining with 200 nM Van-BODIPY led to substantial background fluorescence (Figure 3D), whereas staining of the same tissue with 130 nM Van-JF_669_ produced less background emission (Figure 3E). Moreover, Van-JF_669_ is compatible with antibody labeling, making it appropriate for multicolor experiments in which more than one target is visualized simultaneously (Figure 3F). These experiments confirm the suitability of Van-JF_669_ for detecting *S. aureus* in patient-derived biopsies.

Finally, we hypothesized that the spirocyclization equilibrium that drives the fluorogenicity of Van-JF_669_ could lead to fluorescence blinking,^20^ which could be leveraged to perform single-molecule localization microscopy (SMLM).^21^ We first confirmed that single-molecule blinking could be detected in total-internal reflection fluorescence (TIRF) microscopy (Supporting Video 2). This blinking likely arises from a combination of spontaneous interconversion between the spirocyclic (dark) and zwitterionic (fluorescent) isomers of the JF_669_ dye and transient binding of vancomycin to the peptidoglycan layer of bacteria. The spontaneous blinking of probe Van-JF_669_ allowed us to perform live-cell SMLM by simply incubating *S. epidermidis* with Van-JF_669_ (0.1 nM) without the need for cell fixation or the use of special blinking buffers. SMLM of live *S. epidermidis* (Figure 4A) produced images that had nearly one order of magnitude better resolution than confocal images. For example, the cell wall of *S. epidermidis* in confocal imaging appeared to have a width of ∼260 nm, whereas SMLM using Van-JF_669_ gave images in which the cell-wall has a width of ∼40 nm (Figure 4b), which is consistent with direct observations for Gram-positive bacteria by cryo-electron microscopy.^22^ These results demonstrate that Van-JF_669_ can be used for live-cell super-resolution imaging of the cell wall of *Staphylococcus* bacteria.

**Figure 4.**
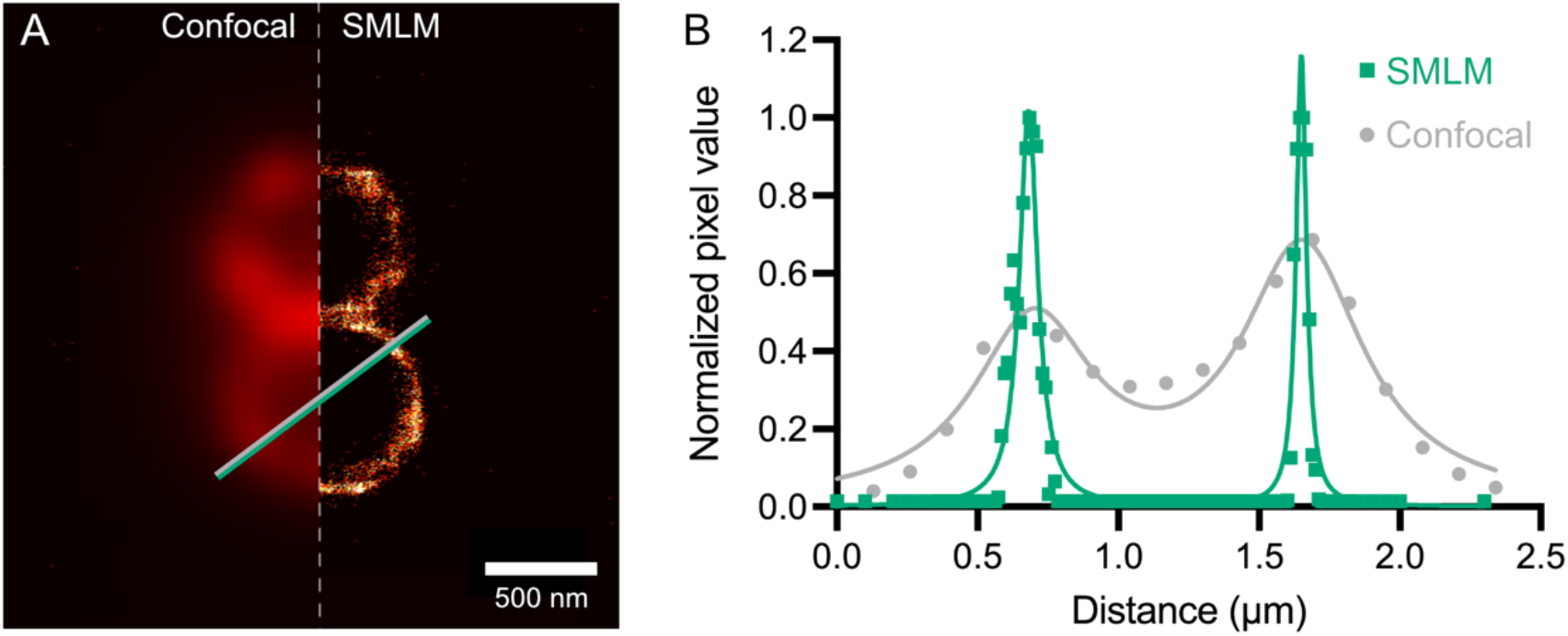
Super-resolution imaging of *S. epidermidis* cell wall. (A) Confocal micrograph (left) and SMLM (right) images of *S. epidermidis* treated with 0.1 nM Van-JF_669_ and imaged in ultrapure water. (B) Pixel values and Lorenzian fitting of these values along the line displayed in panel (A).

## Discussion

In summary, we developed a far-red fluorescent probe to detect staphylococci (and potentially other Gram-positive bacteria) in in vitro bacterial culture, cell-culture infections, and human tissues from patients with deep-seated infections. This probe is easy to synthesize, it can be used to stain complex samples, does not rely on genetic manipulation of the specimen, has virtually no background fluorescence, and is compatible with flow cytometry, fluorescence microscopy and live-cell, super-resolution microscopy. Thus, Van-JF_669_ is a convenient and reliable tool to visualize *Staphylococcus* bacteria in complex samples.

## Methods

### Immunohistochemistry of vibratome sections of human biopsies

60 μm vibratome sections from PROSA329 (stored in cryoprotectant buffer at - 20°C) were washed three times for 5 min with Tris-buffered saline pH 7.4 (TBS) at room temperature.^23^ For antigen retrieval, sections were incubated in 1 mL prewarmed 10 mM sodium citrate, pH 8.5, at 37 °C for 30 min. Sections were washed three times with TBST (0.1% Triton X-100 in TBS), blocked for 30 min with 1% BSA-fraction V, 2% human serum in TBST, and washed again three times with TBST. Endogenous biotin sites were blocked using Streptavidin/Biotin blocking kit (Vector Laboratories, VC-SP-2002) following manufacturer’s protocols. Sections were incubated in 10 μg mL^−1^ anti-*Staphylococcus aureus* antibody (Abcam, ab20920) in blocking buffer (with 1% BSA-fraction V, 2% human serum in TBST) at 4 °C overnight, washed three times with TBST, and stained with 5 μg mL^−1^ Alexa Fluor 568 donkey anti-rabbit IgG (Invitrogen, A10042) for 1 h at room temperature. Sections were washed three times with TBS and incubated with Van-JF_669_ (130 nM) and DAPI (3 μM) for 10 min. Sections were washed three times with TBS, mounted in 40 μL of mounting medium (Agilent DAKO, Cat No. S3023) and examined using an Olympus SpinSR Spinning disc microscope.

### Flow cytometry

Co-cultures were performed with a multiplicity of infection (MOI) of 2. A clinical isolate of *S. aureus* (PROSA25) (10^6^ cells from an overnight culture resuspendend in RPMI + 10% FBS) and THP1 cells (5 × 10^5^ cells in RPMI + 10% FBS) were co-cultured in a 96 well-plate at 37 °C with 5% CO_2_ for 2 h. Afterwards, the plates were centrifuged for 5 min at 350 × G. Each well was blocked using 50 μL of FACS buffer (PBS + 5% FBS) + 2.5 μL Fc receptor (BioLegend, Cat No. 422302) block per well and then 150 μL FACS buffer was added and spun down. The sample was fixed by addition of 5 μL of 4% paraformaldehyde for 20 min in the dark at room temperature. After the addition of 150 μL of permeabilization buffer (BioLegend, Cat No. 421002), the sample was centrifuged. Subsequently, Van-JF_669_ (1.2 μM) and DAPI (3 μM) in permeabilization buffer were stained for 15 min in the dark at room temperature. 150 μL of FACS buffer was added. After spinning down and resuspending in 200 μL PBS, samples were analyzed by flow cytometry (Cytoflex) and image stream.

### Staining of FFPE sections

Formalin-fixed paraffin-embedded patient biopsies were cut into 4 μm-thick sections by microtome (Leica, SM2010R). Samples were deparaffinized by washing twice with for 10 min. Rehydration was carried out by incubating twice with 100% ethanol for 5 min, 96% ethanol for 2 min, 70% ethanol for 2 min, 50% ethanol for 2 min, and distilled water for 2 min. Sections were incubated with EDTA based BOND Epitope Retrieval solution, pH = 9.0 (Biosystems, AR9640) at 95 °C for 20 min for antigen retrieval. After cooling for 15 min, sections were transferred in PBST for 5 min. The slide was dried with Kimwipes and circle was drawn around the sections with a hydrophobic pen. 300 μL of blocking buffer (5% goat serum in PBS + 0.1% Tween 20 (PBST) was added to slides and the slide was incubated in a dark humidity chamber for 1 h at room temperature. After removing the blocking buffer, the primary rabbit anti human anti-Rab5 antibody (Abcam, ab218624) was incubated overnight at 4 °C in a dark humidity chamber. Next, the primary antibody was removed and rinsed three times with PBST for 5 min. Samples were incubated with a secondary antibody at room temperature in a dark humidity chamber for 1 h. Subsequently, the secondary antibody was removed and rinsed three times with PBST for 5 min. Van-JF_669_ (130 nM) or Van-BODIPY (200 nM) and DAPI (3 μM) were incubated at room temperature in a dark humidity chamber for 10 min. After rinsing three times with PBS for 5 min, the slide was mounted and visualized by confocal microscopy.

### Tissue collection

Fresh tissue samples were collected intraoperatively by an orthopedic surgeon at the site of clinical apparent infection from deep-seated *S. aureus* major bone and joint infections. The patients were included and operated between November 2020 and November 2022 at the University Hospital of Basel, Switzerland by orthopedic surgeons specialized in musculoskeletal infections. Fresh samples were either directly used for flow cytometry and or directly fixed in 4% formaldehyde until further use. The study was approved by the local Ethical Review Board (Ethikkommission Nordwestschweiz, Project-ID 2020-02588) and performed in compliance with all relevant ethical regulations.

## Materials availability

Probe Van-JF_699_ is available from the authors at no cost upon reasonable request.

## Data availability

All raw and processed data supporting this manuscript is available on Zenodo. DOI: 10.5281/zenodo.8314732.

## Author contributions

The probe was designed by K. J., D. B., and P. R.-F. K. J. synthesized and characterized Van-JF_669_ in vitro, performed live-cell imaging with *S. epidermidis*, and super-resolution microscopy. V. T. performed MIC determination, probe working concentration determination and validation in patient samples, and *S. aureus* imaging in fixed patient samples and microfluidics. B. M. performed imaging of patient samples and of *S. aureus* co-cultured with macrophages. A. I. performed the co-culture of THP-1 cells with *S. aureus* and contributed to flow cytometry experiments. R. K., M. M., and C.M. developed the clinical sampling workflow and collected the intraoperative patient samples. N. K., D. B., and P. R.-F. supervised the project.

## Declaration of Interests

The authors declare no competing financial interests.

## Acknowledgements

This work was funded by the Swiss National Science Foundation through the National Center of Competence in Research AntiResist (grant 180541). B. M. was additionally funded by the MD-PhD Scholarship from the Swiss National Science Foundation and the Swiss Academy of Medical Sciences (SNF 323530_207037). V. T. was funded by a Biozentrum PhD fellowship. We thank Dr. Gina Grammbitter for assistance with mass spectrometry experiments. We thank the microscopy core facility, flow cytometry facility at the Department of Biomedicine and imaging core facility at Biozentrum, University of Basel.

## Notes

### Competing Interest Statement

The authors have declared no competing interest.

