## Supplementary Information for "A Far-Red Fluorescent Probe to Visualize *Staphylococcus aureus* in Patient Samples"

#### Contents

1 GENERAL REMARKS 2

2 MATERIALS AND METHODS 2

3 SUPPLEMENTARY FIGURES 4

4 SYNTHETIC PROCEDURES 7

#### 1 General remarks

All reagents and solvents were purchased from commercial sources and used as received. NMR spectra were acquired on Bruker AVANCE NEO-400, Bruker AVANCE III-400, Bruker AVANCE III HD-600, and Bruker AVANCE II-800 instruments. $^{1}$H NMR chemical shifts are reported in ppm relative to $SiMe_4$ ($\delta$ = 0) and were referenced internally with respect to residual protons in the solvent ($\delta$ = 1.94 for acetonitrile and $\delta$ = 3.31 for methanol). Coupling constants are reported in Hz. $^{13}$C NMR chemical shifts are reported in ppm relative to $SiMe_4$ ($\delta$ = 0) and were referenced internally with respect to solvent signal ($\delta$ = 1.32 for acetonitrile and $\delta$ = 49.00 for methanol). High-resolution mass spectrometry (HRMS) was performed by the MS facility of EPFL and UZH. Reaction progress was followed by ultra-high performance liquid chromatography–mass spectrometry (UHPLC-MS) on a Shimadzu LC-MS 2020 system using electrospray ionization (ESI) and Low-resolution mass spectra (LRMS) were acquired on a Waters spectrometer by using electrospray ionization (ESI). Purification by prep-HPLC was performed using a Büchi Pure-Chromatography-System and Büchi FlashPure columns. IUPAC names of all compounds are provided and were determined using CS ChemDraw 20.1.

#### 2 Materials and methods

**Optical spectroscopic methods**
Stock solutions were prepared in DMSO (spectrophotometric grade > 99.9%) at concentrations of 5 mM and stored at -80°C. Spectroscopic measurements were conducted in phosphate-buffered saline (PBS). UV-Vis spectra were acquired using a Multiskan SkyHigh Microplate Spectrophotometer (ThermoFisher Scientific) and quartz cuvettes from Thorlabs (10 mm path length) or 96-well plates (Corning). Measurements were carried out as stated at 25°C unless stated otherwise. Fluorescence spectra were acquired using a FS5 Spectrofluorometer (Edinburgh Instruments) equipped with a SC-25 cuvette holder or SC-40 plate reader and quartz cuvettes from Thorlabs (10 mm path length) or 96-well plates (Corning). All measurements were carried out as three technical replicates. The obtained UV-Vis spectra were background corrected. Prism 8.0 was used to process the data and plot the spectra.

**Bacterial culture**

Bacteria, *S. epidermidis* (ATCC 12228) was grown in TSB in 5 mL culture tubes at 37°C, shaking (180 rpm). Clinical isolated *S. aureus* (PROSA28), lab strain MSSA (ATCC 29213), and lab strain MRSA (ATCC 43300) were grown in 5 mL cation-adjusted MHB in 50 mL culture tubes at 37°C, shaking (180 rpm). Overnight cultures were streaked on TSB agar plates at 37°C for 16-18 hours. A single colony was picked and inoculated in TSB/cation-adjusted MHB in 5 mL culture tubes at 37°C, shaking (180 rpm) for 16-18 hours.

**Live-cell microscopy**

All incubations were done at 37°C and cultures were shaken at 180 rpm. We transformed PROSA with a pRN11-derivative[1] carrying a transcriptional fusion of the $P_{pdhABCD}$ promoter and gfp-mut3.1 obtained from plasmid pC183-S3[2] resulting strain called PROSA28-pVT13. PROSA28-pVT13 was streaked on agar containing brain-heart infusion (BHI) (Remel, Cat No. 452472) with 10 mg $L^{-1}$ chloramphenicol. Overnight cultures were prepared by picking a single colony and inoculating BHI containing 10 mg $L^{-1}$ chloramphenicol. Overnight cultures were diluted 1:1000 and grown for 2 hours. $OD_{600}$ was adjusted to 0.003 in BHI and loaded in a CellAsic plate (Millipore, Cat No. B04A-03). Bacteria were flushed with BHI + 0.17 μM unlabeled vancomycin + 0.13 μM Van-$JF_{669}$ for 150 minutes at 10 kPa. Images were taken every two minutes using a Nikon Ti2 widefield microscope with 100X objective (Laser powers 5% in ch640 and 20% in ch488, exposure times 200 ms)

**Super-resolution microscopy**

A μ-slide, 8-well, glass-bottom plate (Ibidi, Cat No. 80827) was cleaned with UV-generated ozone for 10 minutes and held for 5 minutes at 25 °C. A *S. epidermidis* suspension in LB broth ($OD_{600}$=0.1) was added to the well and incubated at 25 °C for 10 minutes. After washing three times of milliQ water, 0.1 nM Van-$JF_{669}$ in water was added to the well. Images were taken using a combined Nikon W1 spinning disc (confocal) and N-STORM (SMLM) system equipped with a Nikon 1.49 NA 100x TIRF Apo Plan SR objective. After optimization of the HILO angle, the acquisition of frames was started, using a 638 nm laser at 90 mW power, with an exposure time of 30 ms and a constant frame rate (15.5 Hz). The acquisition was continued until 5000 frames were required. Images were processed by the Picasso software package (https://github.com/jungmannlab/picasso).[3]

### 3 Supplementary figures

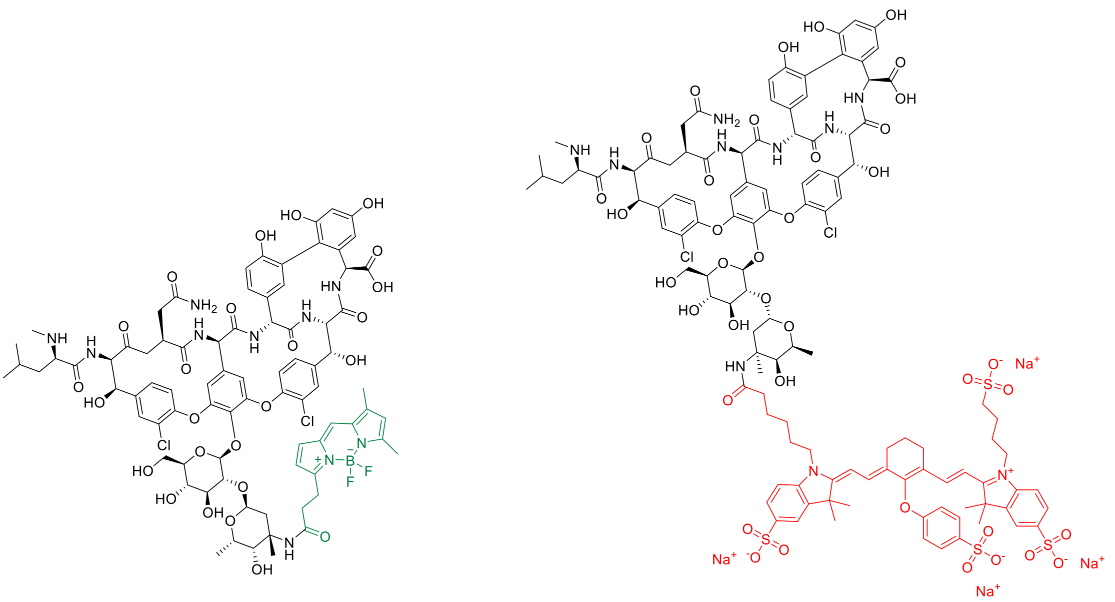

**BODIPY™ FL Vancomycin (Van-BODIPY)** **Vanco-800CW**

**Figure S1.** Chemical structure of Van-BODIPY and Vanco-800CW.

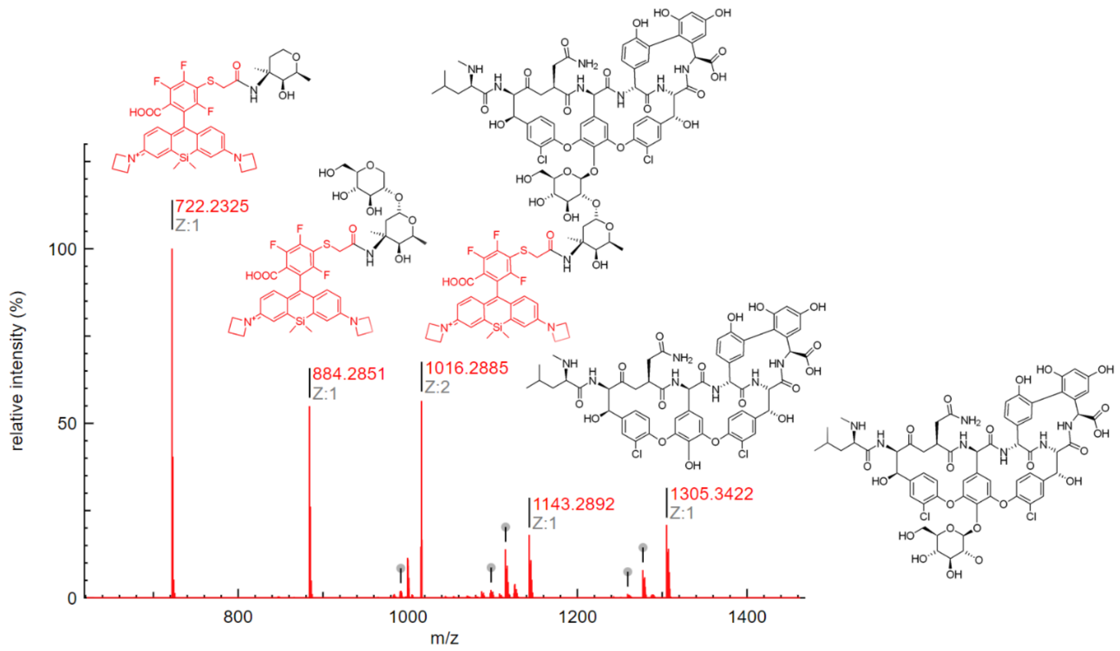


**Figure S2.** Structure determination of Van-JF$_{669}$. The fragments detected by HRMS correspond to JF$_{669}$ attached to the sugar moiety of vancomycin.

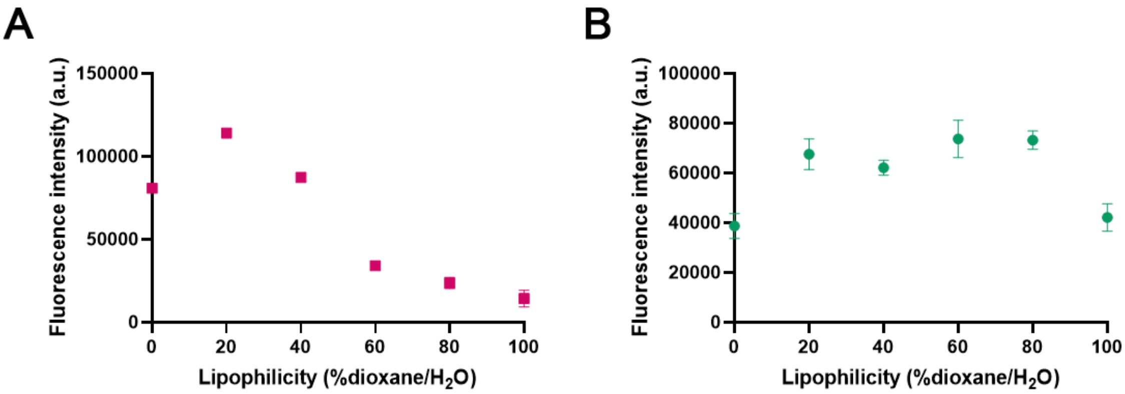


**Figure S3.** Fluorescence of vancomycin conjugated to different fluorophores. (A) Lipophilicity-dependent changes of Van-JF669 fluorescence and (B) lipophilicity-independent changes of Van-BODIPY fluorescence. Symbols indicate mean and whiskers the standard deviation from three technical replicates.

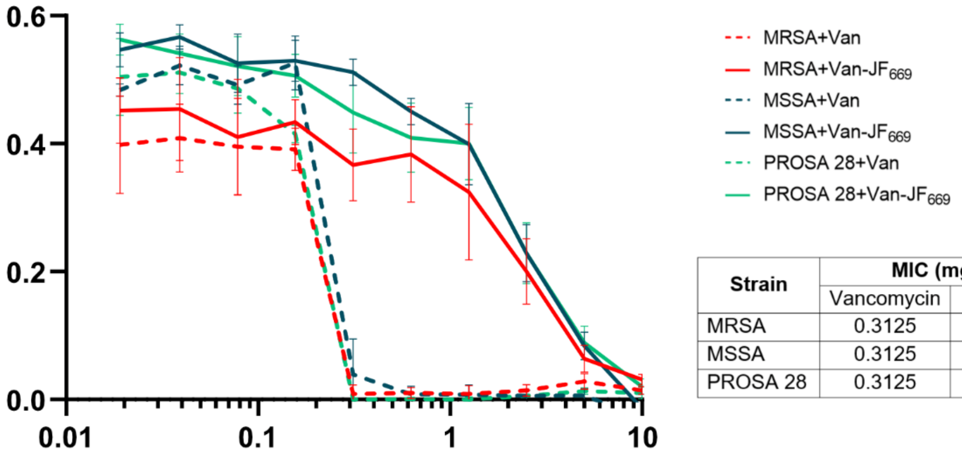


| Strain | MIC (mg L$^{-1}$) | |
|---|---|---|
| | Vancomycin | Van-$JF_{669}$ |
| MRSA | 0.3125 | 5 |
| MSSA | 0.3125 | 5 |
| PROSA 28 | 0.3125 | 5 |

**Figure S4.** The minimum inhibitory concentration of vancomycin(van) and Van-JF669 in methicillin-resistant *S. aureus* (MRSA, ATCC43300), methicillin-sensitive *S. aureus* (MSSA, ATCC 29213), and clinical isolate *S. aureus* (PROSA28, methicillin sensitive). Lines indicate mean and whiskers standard deviation from four biological replicates.

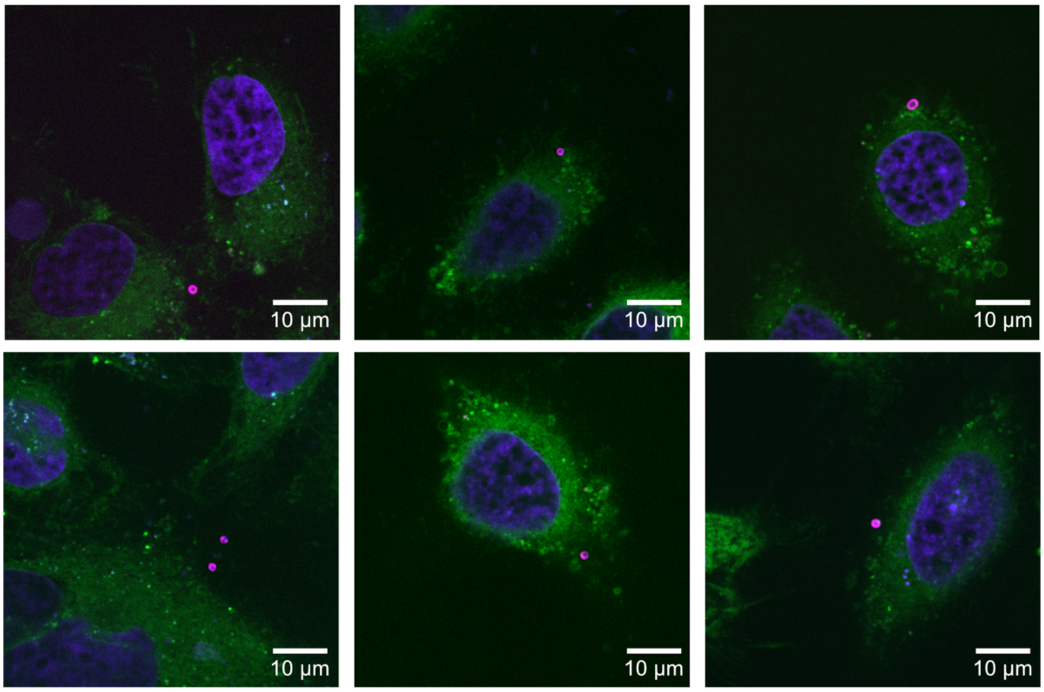


**Figure S5**. Further examples of micrographs showing selective labeling of *S. epidermidis* co-cultured with HeLa cells and treated with 1.5 μM Hoechst (nuclei = blue), 2 μM ER-Tracker™ Green (endoplasmic reticulum = green) and 10 nM Van-JF$_{669}$ (*S. epidermidis* = magenta).

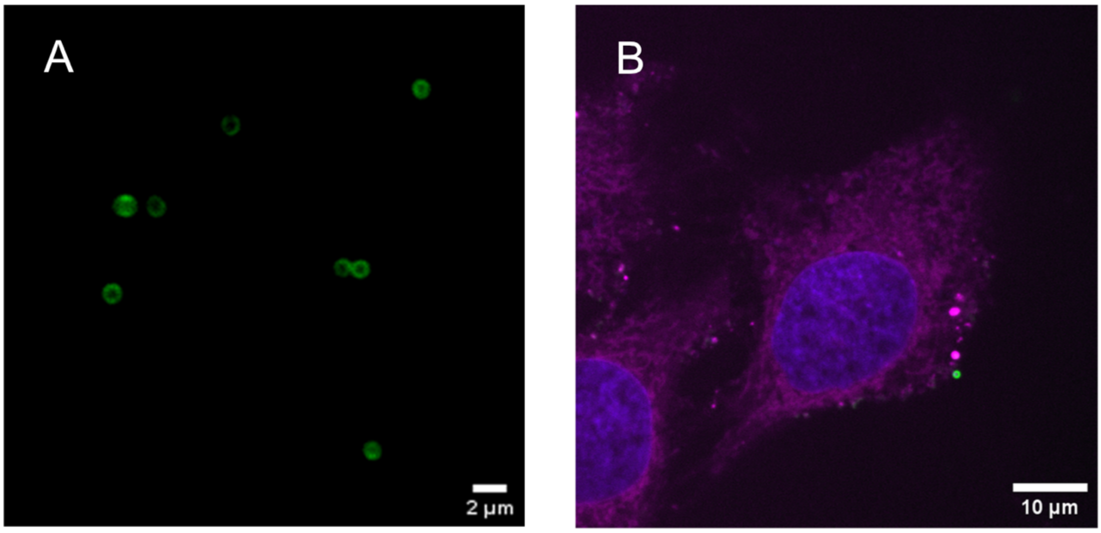


**Figure S6.** Fluorescent images *S. epidermidis* labeled with Van-BODIPY; (A) *S. epidermidis* was treated with 2 nM of Van-BODIPY, (B) *S. epidermidis* co-cultured with HeLa cell were treated with 1.5 μM Hoechst (nuclei = blue), 1 μM ER-Tracker™ Red (ER = magenta) and 10 nM Van-BODIPY (*S. epidermidis* = green).

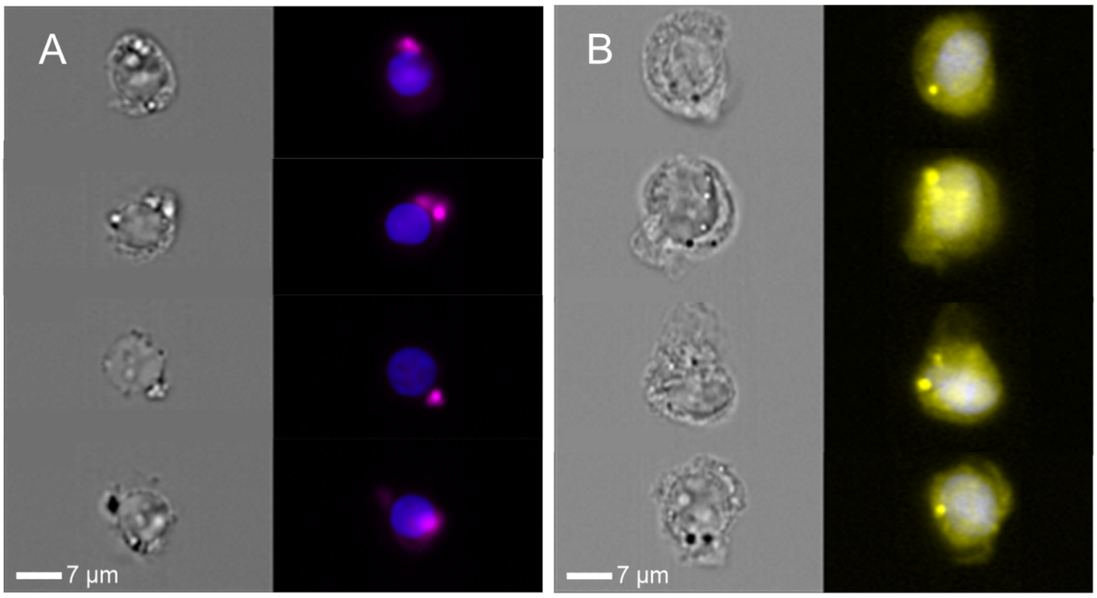


**Figure S7.** Further examples of flow cytometry images of in vitro co-culture of THP1 cells + clinical *S. aureus* (PROSA25) labeled by (A) 1.2 µM Van-$JF_{669}$ and (B) 4.84 µM RADA treated with 3 µM DAPI (nuclei = blue).

### 4 Synthetic procedures

**Scheme S1.** Synthesis of vancomycin containing $JF_{669}$ (Van-$JF_{669}$)

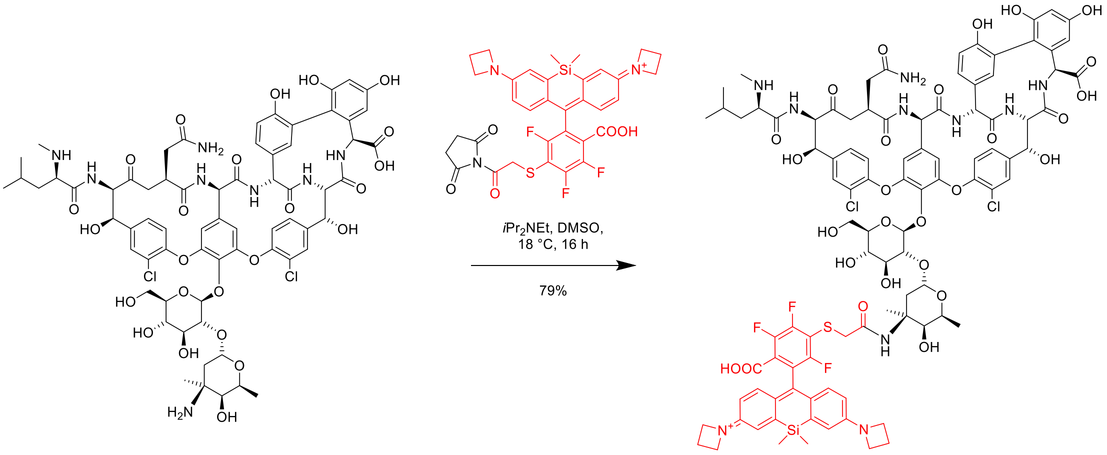

1-(10-(3-((2-(((2S,3S,4S,6S)-6-(((2S,3R,4S,5S,6R)-2-(((3S,6R,7R,22R,23S,26S,36R,38aR)-3-(2-amino-2-oxoethyl)-26-carboxy-10,19-dichloro-7,22,28,30,32-pentahydroxy-6-((R)-4-methyl-2-(methylamino)pentanamido)-2,5,24,38,39-pentaoxo-2,3,4,5,6,7,23,24,25,26,36,37,38,38a-tetradecahydro-1H,22H-23,36-(epiminomethano)-8,11:18,21-dietheno-13,16:31,35-di(metheno)benzo[n][1]oxa[6,9]diazacyclohexadecino[4,5-d][1]oxa[7,17]diazacyclotetracosin-44-yl)oxy)-4,5-dihydroxy-6-(hydroxymethyl)tetrahydro-2H-pyran-3-yl)oxy)-3-hydroxy-2,4-dimethyltetrahydro-2H-pyran-4-yl)amino)-2-oxoethyl)thio)-6-carboxy-2,4,5-trifluorophenyl)-7-(azetidin-1-yl)-5,5-dimethyldibenzo[b,e]silin-3(5H)-ylidene)azetidin-1-ium (**Van-JF$_{669}$**)

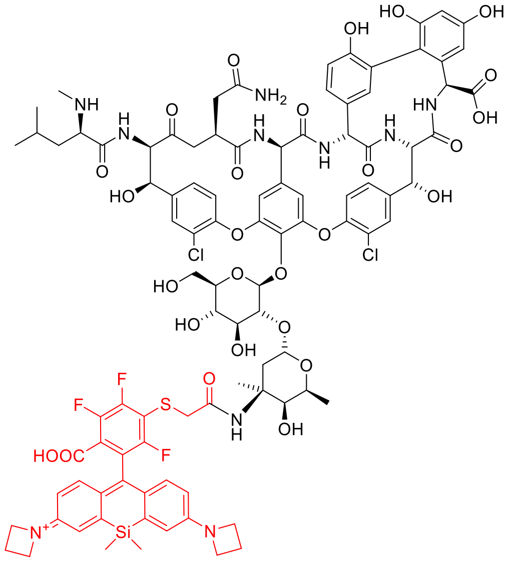

Vancomycin hydrochloride (1 mg, 0.67 μmol,), JaneliaFluor 669 NHS ester (700 μg, 1 μmol) and diisopropylethylamine (1 μL, 6 μmol) were stirred in dimethyl sulfoxide (500 μL) at room temperature for 16 hours. After that, the reaction was lyophilized. Next, acetonitrile (1 mL) and water (1 mL) were added and then purified by semi-preparative HPLC (elution gradient from acetonitrile/water = 1:9 to acetonitrile/water = 7:3) to give a blue powder (1 mg, 79%)

HRMS (ESI) $[M+H]^{2+}$ calculated for $[C_{96}H_{101}Cl_2F_3N_{11}O_{27}SSi]^+$: 1014.2894, found 1014.2879

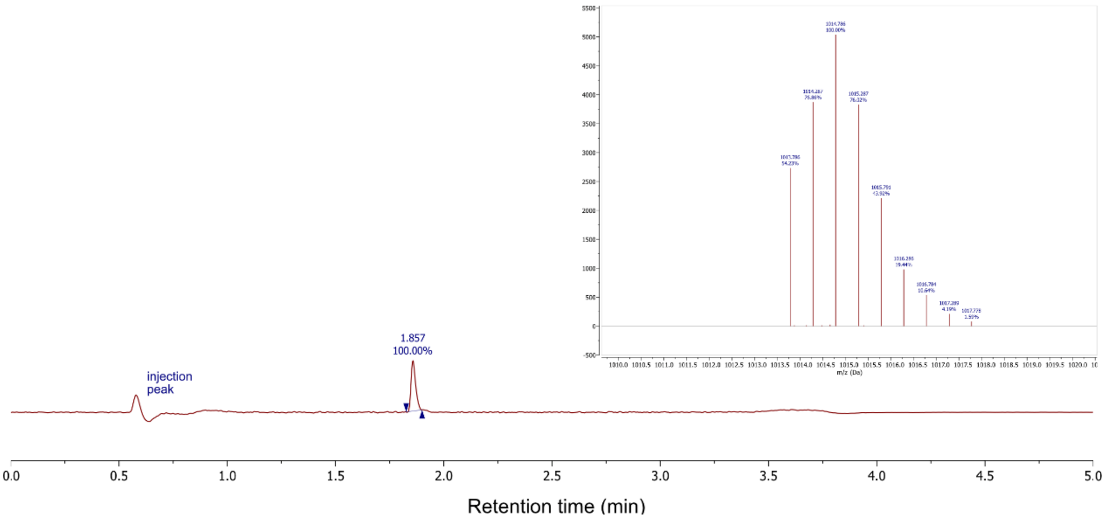


**Figure S8.** Integrated absorbance signal (100-300 nm) of analytical LC-MS run (10 → 95% acetonitrile in dd$H_2O$ + 0.02% TFA + 0.04% FA over 3 min, then additional 2 min at 95%) of Van-$JF_{669}$.
